# A novel B.1.1.523 SARS-CoV-2 variant that combines many spike mutations linked to immune evasion with current variants of concern

**DOI:** 10.1101/2021.09.16.460616

**Authors:** Brian M.J.W. van der Veer, Jozef Dingemans, Lieke B. van Alphen, Christian J.P.A. Hoebe, Paul H.M. Savelkoul

## Abstract

In the severe acute respiratory syndrome coronavirus 2 (SARS-CoV-2) pandemic several variants have emerged that are linked to increased transmissibility and immune evasion. These variants are recognized as variants of concern (VOC). In this study, we describe a B.1.1.523 variant that shares many spike mutations with current VOC. Receptor-binding domain mutations E484K and S494P were observed but also a deletion (position 156-158) in the N-terminal antigenic supersite that is similar to the delta-variant. These mutations are linked to immune evasion in VOC that could lead to less effective vaccines. This variant has been reported in various different countries and continents despite the dominance of B.1.1.7 (alpha) and B.1.617.2 (delta) variant. Furthermore, the B.1.1.523 pangolin lineage as a whole is recognized as a variant under monitoring since 14^th^ of July 2021.

## Introduction

Severe acute respiratory syndrome coronavirus 2 (SARS-CoV-2) infected millions of people during the pandemic ^1,2^. During this pandemic, various variants of this virus were detected in surveillance and were linked with increased infectivity or immune evasion ^3,4^. Data of these variants are shared via the GISAID (Global initiative on sharing all influenza data) database that helps to understand spread and evolution of SARS-CoV-2. The Centers for Disease Control and Prevention (CDC) and European Centre for Disease Prevention and Control (ECDC) assign certain variants as “variant of concern” (VOC) because of an increase in transmissibility, more severe disease, or immune evasion ^5,6^. Current VOC are B.1.1.7 (alpha-variant), B.1.351 (beta-variant), P1 (gamma-variant), and B.1.617.2 (delta-variant). These VOC harbor several spike protein mutations linked to immune evasion. The receptor-binding domain (RBD) and N-terminal domain (NTD) are frequently targeted by neutralizing antibodies mostly directed to the NTD ^7,8^. In addition, a so called antigenic supersite is described in the NTD with three regions. Potent neutralizing antibodies in convalescent plasma target this antigenic super site. Still, mutations in the RBD are important as for example E484K mutation is also strongly linked with immune evasion ^3,4^. This short communication describes a new variant with a new combination of various concerning spike mutations shared with VOCs and is already spread across many countries. The pangolin lineage of this variant is B.1.1.523 and is recognized as a variant under monitoring since 14^th^ of July 2021 ^9^.

## Methods

### GISAID data download and SARS-CoV-2 sequencing

All sequences and metadata of B.1.1.523 variant cases with spike mutations S:E156del, S:F157del, S:R158del, S:E484K, and S:S494P were downloaded from www.gisaid.org on 19 August 2021 (n=551). These mutations were chosen because of links with immune evasion. Of these cases 18 were removed based on bad quality score in Nextclade (clades.nextstrain.org). In routine SARS-CoV-2 surveillance one case of this B.1.1.523 variant was identified with nanopore sequencing as described in von Wintersdorff et al. 2021 ^10^.

### Data-analysis

Visualization of the filtered data by country, month, and age group were made in R statistical software, version 3.6.2 (R Foundation for Statistical Computing, Vienna, Austria). Figures were made using the R package ggplot2 version 3.3.2. Phylogenetic analysis was performed as described in von Wintersdorff et al. 2021 but with nextstrain/ncov version 7 (https://github.com/nextstrain/ncov) ^10^. The amino acid sequence of Wuhan strain, VOC, and B.1.1.523 variant were aligned in MEGA v10.0.5 with Cluster Omega algorithm, with the Wuhan strain as a reference. The 3D structure of the spike protein of the B.1.1.523 variant was predicted using CoVSurver (https://www.gisaid.org/epiflu-applications/covsurver-mutations-app/)

## Results

In total, 533 cases of B.1.1.523 with spike mutations S:E156del, S:F157del, S:R158del, S:E484K, and S494P are reported in GISAID till 19 August 2021. Most cases are reported in Russia followed by Germany but cases have also been seen in the USA and Australia (figure 1A). The first few cases were collected in February 2021 and increased in numbers to 203 cases in May 2021 (figure 1B). Number of cases is lower in June and July 2021 as sequenced-based surveillance data is typically lagging. This variant did not show an age related pattern (figure 1C).

**Figure 1.**
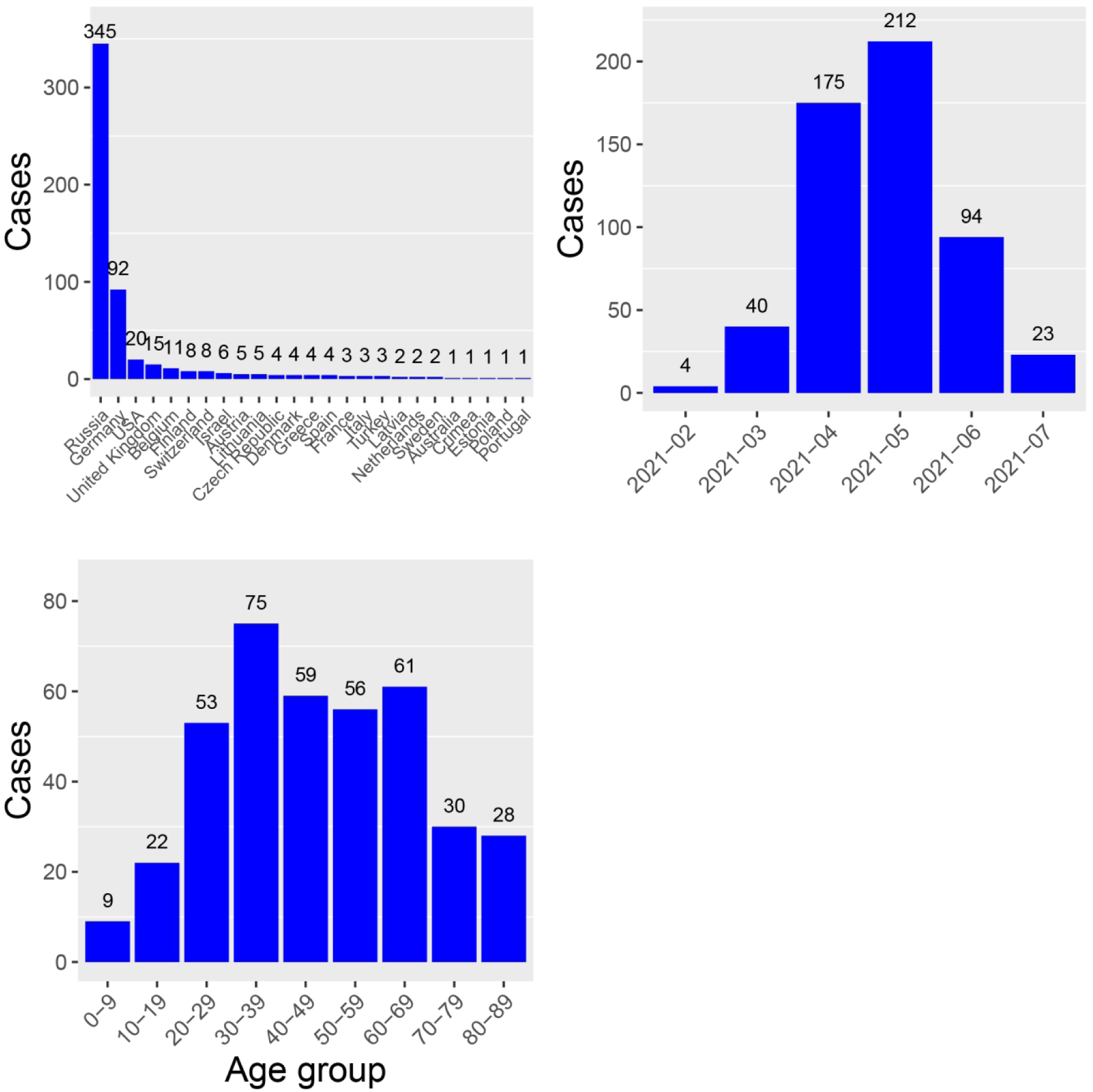
(A) Number of cases per country. (B) number of cases per month, note that data of June and July are incomplete as surveillance is typically lagging. (C) Cases per age group.

Sharing of the pangolin lineage and some spike mutations does not necessarily imply a single origin of this variant. Also, recognition of the first cases in Russia does necessarily implicates that this variant originates from this country. To address both issues a phylogenetic tree was constructed and showed that all cases are similar as they are in the same branch (figure 2). Based on this phylogenetic analysis the origin of this variant is likely Russian as the first strain is reported from Moscow (green circle at 30 mutations (EPI_ISL_1823183), figure 2).

**Figure 2.**
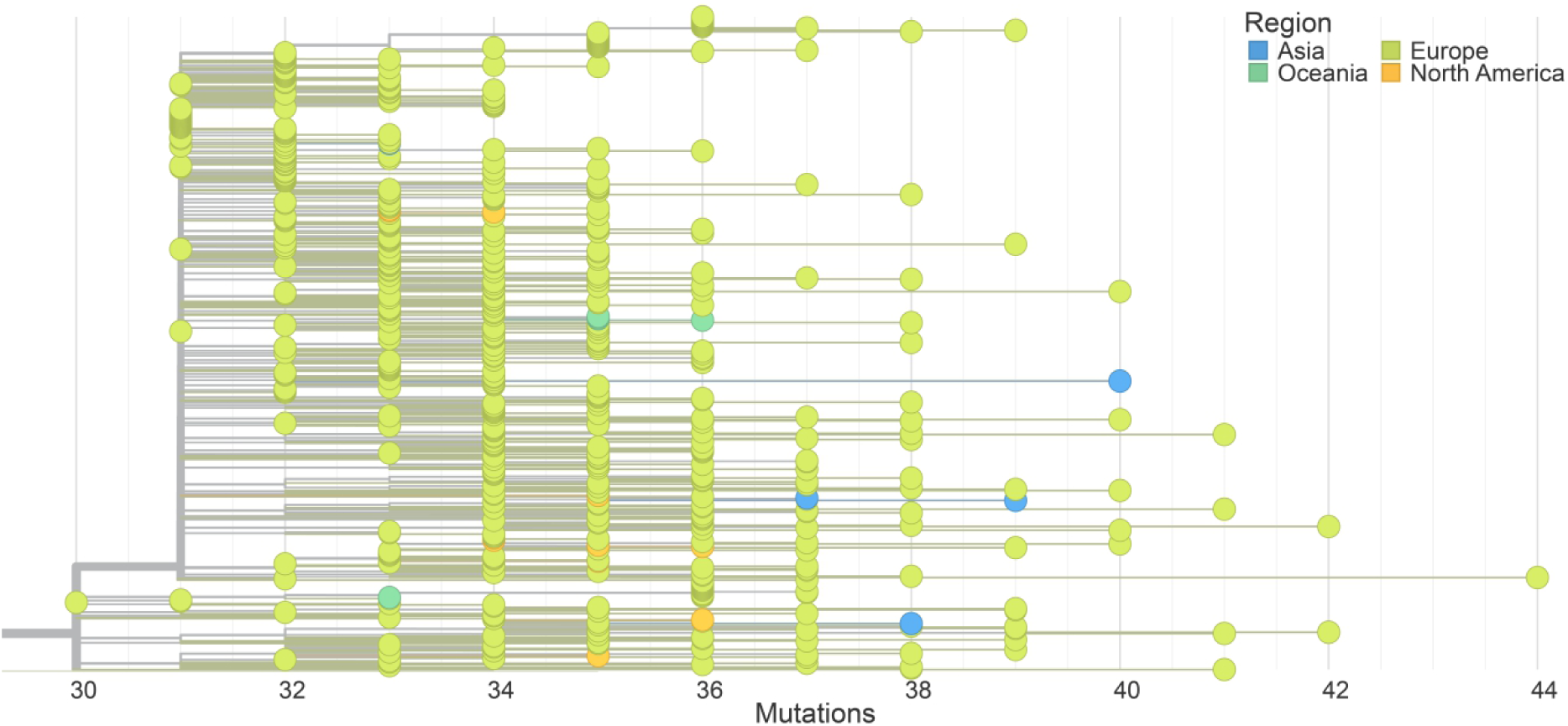
Phylogenetic analysis of cases of B.1.1.523 lineage with S:E156del, S:F157del, S:R158del, S:E484K, and S:S494P.

One of the reasons of concern about this variant is a three amino acid deletion in the NTD antigenic supersite and E484K mutation of the spike protein. Therefore a multiple sequence alignment (MSA) was performed with the amino acid sequence of VOC and Wuhan-Hu-1 (figure 3A). Three VOC, B.1.1.7, B.1.351, and B.1.617.2, have deletions in one of the regions of the NTD antigenic supersite. The deletion of B.1.1.523 is similar to the one of B.1.617.2 and has the E484K mutation that is shared in many VOC. In figure 3B the predicted spike structure and interaction with human ACE2 receptor is shown. Spike mutations E484K and S494P are both in the RBD with the ACE2 receptor.

**Figure 3.**
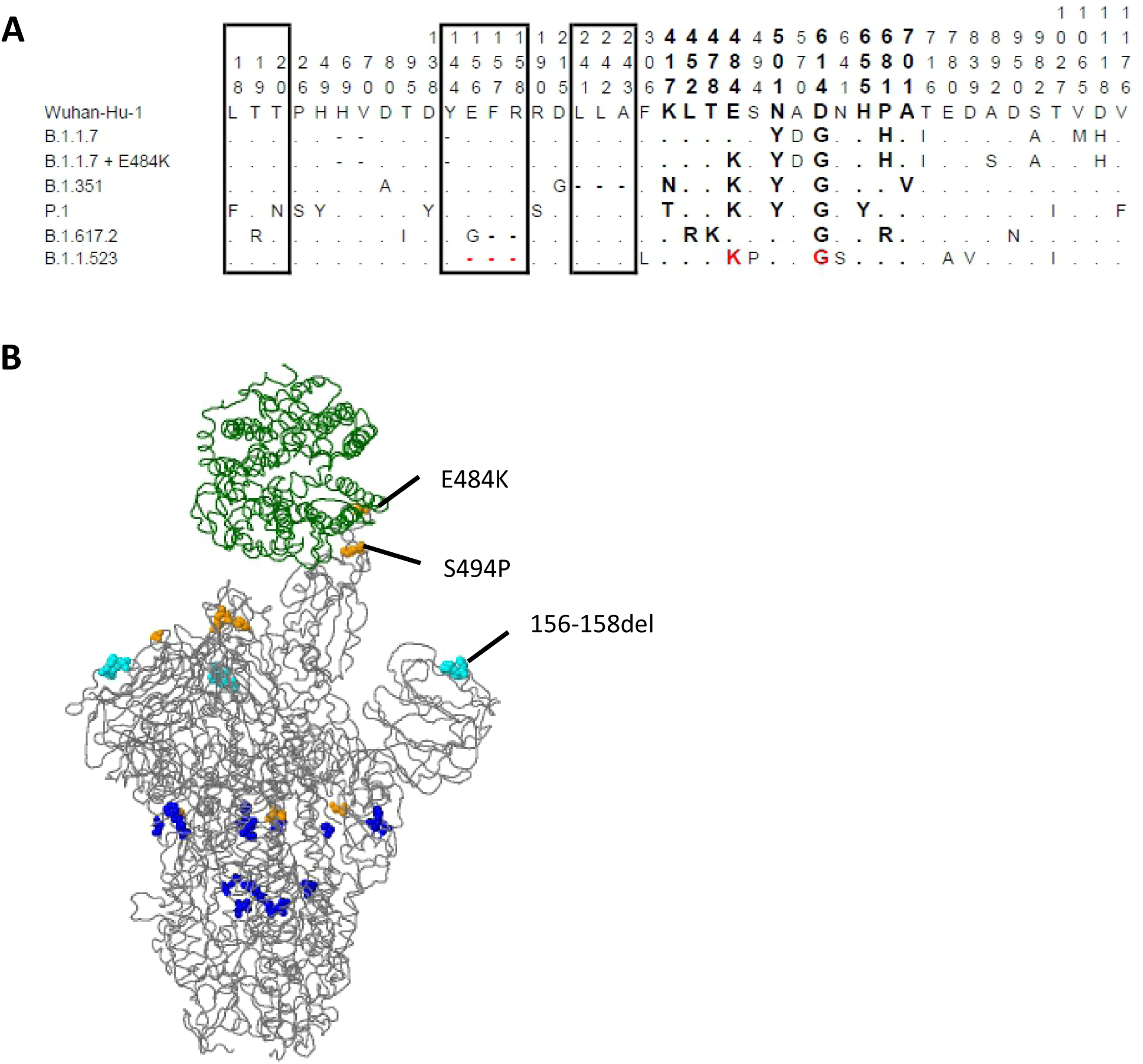
(A) Multiple sequence alignment of the spike protein of Wuhan strain, variants of concern, and B.1.1.523 lineage. In bold are spike mutations linked to increased infectivity or immune evasion in variants of concern. In red are the mutations (or position) in B.1.1.523 that is shared with variants of concern. The block boxes represent mutations in the antigenic supersite of the N-terminal domain of the spike protein. (B) Predicted 3D-structure of the spike protein of the B.1.1.523 lineage using CoVSurver.

## Discussion

In this short communication we report a variant (B.1.1.523) with a new combination of concerning spike mutations that are shared with current VOCs. Many of these mutations are linked with immune evasion that could lead to less effective vaccines. This variant has been reported in various different countries and continents and likely originates in Russia. In addition, the number of cases appears to increase despite the dominant variants B.1.1.7 (alpha) and B.1.617.2 (delta).

Mutations that are shared with VOC or linked with immune evasion were S:E156del, S:F157del, S:R158del, S:E484K, and S:S494P ^5,6,11^. Two mutations are in the RBD of the spike protein, positions 484 and 494, and are both linked to immune evasion in B.1.1.7 (alpha) variant. E484K mutation is also present B.1.351 (beta) and P.1 (gamma) variants that are strongly linked with reduced efficacy of vaccines ^3,4^. These findings are supported in studies where the effect of spike mutations on efficacy of monoclonal antibodies and convalescent plasma was investigated ^12^. Similar studies were performed to investigate the antigenic super site of the NTD ^7,8^. In the β-hairpin region of the antigenic super site the B.1.1.523 variant has a deletion (position 156-158) similar to the currently dominant B.1.617.2 (delta) variant (S:E156G and 157-158del) ^5-8^. Other VOC also have deletions in one of the regions of the antigenic super site, B.1.1.7 (alpha-variant) position 144 and B.1.351 (beta-variant) position 241-243. As the mutations observed in this B.1.1.523 variant are strongly linked to immune evasion and disseminated to different continents, despite the dominance of both the B.1.1.7 (alpha) and B.1.617.2 (delta) variants, this could be a variant of interest and should be monitored closely. However, as this is the first study that describes this variant of the B.1.1.523 lineage, there is yet no information on transmissibility that contributes to the need of actions required to prevent dissemination.

## References

1 Li, Q. et al. Early Transmission Dynamics in Wuhan, China, of Novel Coronavirus-Infected Pneumonia. N Engl J Med 382, 1199–1207, doi:10.1056/NEJMoa2001316 (2020).

2 WHO. Coronavirus (COVID-19) Dashboard, < https://covid19.who.int/ (2021).

3 Peacock, T. P., Penrice-Randal, R., Hiscox, J. A. & Barclay, W. S. SARS-CoV-2 one year on: evidence for ongoing viral adaptation. J Gen Virol 102, doi:10.1099/jgv.0.001584 (2021).

4 Harvey, W. T. et al. SARS-CoV-2 variants, spike mutations and immune escape. Nat Rev Microbiol, doi:10.1038/s41579-021-00573-0 (2021).

5 Center for Disease Control and Prevention. SARS-CoV-2 Variant Classifications and Definitions. https://www.cdc.gov/coronavirus/2019-ncov/variants/variant-info.html assessed 27 August 2021

6 European Centre for Disease Prevention and Control. SARS-CoV-2 variants of concern. https://www.ecdc.europa.eu/en/covid-19/variants-concern assessed 27 August 2021

7 McCallum, M. et al. N-terminal domain antigenic mapping reveals a site of vulnerability for SARS-CoV-2. Cell 184, 2332–2347 e2316, doi:10.1016/j.cell.2021.03.028 (2021).

8 Suryadevara, N. et al. Neutralizing and protective human monoclonal antibodies recognizing the N-terminal domain of the SARS-CoV-2 spike protein. Cell 184, 2316–2331 e2315, doi:10.1016/j.cell.2021.03.029 (2021).

9 World Health Organisation. Tracking SARS-CoV-2 variants. https://www.who.int/en/activities/tracking-SARS-CoV-2-variants/ assessed 16 September 2021

10 von Wintersdorff, C.J.H., van Alphen, L.B., Wolffs, P.F.G., van der Veer, B.M.J.W., Hoebe, C.J.P.A., Savelkoul, P.H.M.. Infections caused by the Delta variant (B.1.617.2) of SARS-CoV-2 are associated with increased viral loads compared to infections with the Alpha variant (B.1.1.7) or non-Variants of Concern. Research Square, doi:10.21203/rs.3.rs-777577/v1 (2021).

11 Greaney, A. J. et al. Complete Mapping of Mutations to the SARS-CoV-2 Spike Receptor-Binding Domain that Escape Antibody Recognition. Cell Host Microbe 29, 44–57 e49, doi:10.1016/j.chom.2020.11.007 (2021).

12 Weisblum, Y. et al. Escape from neutralizing antibodies by SARS-CoV-2 spike protein variants. Elife 9, doi:10.7554/eLife.61312 (2020).

